# Chemically-induced Neurite-like Outgrowth Reveals Multicellular Network Function in Patient-derived Glioblastoma Cells

**DOI:** 10.1101/467654

**Authors:** Barbara da Silva, Euan S. Polson, Alastair Droop, Ryan K. Mathew, Lucy F. Stead, Jennifer Williams, Susan C. Short, Margherita Scarcia, Georgia Mavria, Heiko Wurdak

**Affiliations:** School of Medicine, University of Leeds, Leeds, LS2 9JT, UK; Leeds Institute for Data Analytics, University of Leeds, Leeds, LS2 9JT, UK; Department of Neurosurgery, Leeds General Infirmary, Leeds, LS1 3EX, UK; Faculty of Biology, Medicine and Health, University of Manchester, Manchester M13 9PT, UK

**Keywords:** Brain tumor, glioblastoma multiforme, patient-derived tumor cells, GBM stem-like cells, ROCK inhibition, neurite-like outgrowth, malignant multicellular network, organelle trafficking, radiation resistance

## Abstract

Tumor stem cells and malignant multicellular networks have been separately implicated in the therapeutic resistance of Glioblastoma Multiforme (GBM), the most aggressive type of brain cancer in adults. We show that small molecule inhibition of RHO-associated serine/threonine kinase (ROCKi) significantly promoted the outgrowth of neurite-like cell projections in cultures of heterogeneous patient-derived GBM stem-like cells. These projections formed *de novo*-induced cellular network (iNet) ‘webs’, which regressed after withdrawal of ROCKi. Connected cells within the iNet web exhibited long range calcium signal transmission, and significant lysosomal and mitochondrial trafficking. In contrast to their less-connected vehicle control counterparts, iNet cells remained viable and proliferative after high-dose radiation. These findings demonstrate a link between ROCKi-regulated cell projection dynamics and the formation of radiation-resistant multicellular networks. Our study identifies means to reversibly induce iNet webs *ex vivo*, and may thereby accelerate future studies into the biology of GBM cellular networks.

## INTRODUCTION

GBM is the most frequent malignant primary brain tumor in adults and despite multimodality treatment, prognosis remains poor (Davis 2016; Grossman and Batara 2004). The majority of GBM patients undergo neurosurgical debulking followed by adjuvant chemoradiotherapy (Davis 2016). However, 5-year survival is less than 5% (Ostrom et al. 2014), and in most cases, tumors recur within the margin zone adjacent to the surgical cavity from microscopic residual tumor cells surviving radiation and chemotherapy (Osuka and Van Meir 2017). The cellular heterogeneity of GBM tumors (Patel et al. 2014), the migratory/brain-infiltrative nature of GBM cells (Demuth and Berens 2004), the appearance and maintenance of poorly-differentiated stem cell-like features (Eyler et al. 2011), the cell-to-cell communication via membrane tubes and formation of a malignant network (Winkler and Wick 2018; Osswald et al. 2015; Weil et al. 2017) are all pathological phenomena that have been implicated in GBM therapeutic resistance. However, the mechanisms underlying these dynamic processes, as well as their interplay and common molecular denominators, remain largely elusive.

The cytoskeleton is a key organizer of cell function, spatially arranging cellular content, establishing connections to the external environment, and enabling cellular movement and change of cell shape (Fletcher and Mullins 2010). The RHO-associated serine/threonine kinase (ROCK) family plays a central role in the regulation of actin cytoskeletal dynamics (Sanz-Moreno et al. 2011). ROCK comprises the ROCK1 and ROCK2 proteins, which phosphorylate substrates including LIM domain kinases 1 and 2 (LIMK1 and LIMK2), myosin light chain (MLC) 2, and the myosin-binding subunit of myosin phosphatase (MYPT1), which regulate actin filament stabilization and myosin-driven contraction, respectively (Maekawa et al. 1999; Ohashi et al. 2000; Rath and Olson 2012). Accordingly, ATP-competitive small molecule inhibitors have been used to study the role of ROCK1 and ROCK2 in cancer cells, and chemical pan-ROCK inhibition (hereinafter referred to as ROCKi) has been considered as a potential anti-tumor therapeutic strategy (Rath and Olson 2012). However, ROCKi approaches have resulted in conflicting pro- and anti-tumor effects in various cancer cell types, including GBM cell lines (Zohrabian et al. 2009; Tilson et al. 2015; Salhia et al. 2005). Furthermore, ROCKi can mediate significant cell morphological changes in neural stem cells (Jia et al. 2016), and has been linked to axonal outgrowth and neuroprotective effects (Dergham et al. 2002; Fournier et al. 2003; Fujita and Yamashita 2014).

Based on this body of literature, we hypothesized that ROCKi-dependent cytoskeletal rearrangements may play a role in GBM cell biology. We sought to clarify the effect of ROCKi by examining the cellular phenotype of well-characterized patient-derived GBM cells upon addition of pan-ROCK inhibitors (Rath and Olson 2012). We observed that ROCKi robustly induced the formation and elongation of GBM cell projections that were morphologically akin to neurites. Furthermore, we investigated the impact of these cell projections on cellular connectivity using mathematical and experimental network analyze and examined multicellular network function in regards with a potential for promoting radiation resistance (Winkler and Wick 2018; Osswald et al. 2015; Weil et al. 2017; Rath and Olson 2012).

## RESULTS

### ROCKi leads to neurite-like projection outgrowth in GBM cell models

We phenotypically characterized chemical ROCK inhibition in transcriptionally heterogeneous patient-derived GBM cell lines that represent different parental tumor origins and molecular GBM subtypes (Drubin et al. 1985; Wurdak et al. 2010; Wurdak 2012; King et al. 2017; da Silva et al. 2018; Polson et al. 2018). We used adherent culture conditions, which conserved their stem cell-like characteristics (Wurdak et al. 2010; Wurdak 2012; King et al. 2017; da Silva et al. 2018; Polson et al. 2018). After treatment of five different cell lines with 20 μM ROCK inhibitor Y-27632, time lapse microscopy imaging revealed a pronounced elongation of cellular projections akin to cytokine-stimulated neurite outgrowth (Drubin et al. 1985) (**Fig 1A, Videos S1-10**). We traced 12 randomly-selected cells per vehicle- or Y-27632-treated GBM cell model comparing initial (t_o_) with 24-hour (t_24_) time points. In the presence of Y-27632, all tested GBM cell models displayed a marked increase in the length of cellular projections compared to the vehicle (H_2O_) control (**p≤**0.02642; **Fig. 1B**). Projections stained positive for the neuron-specific neurite markers Class III β-tubulin (TuJ1) and the microtubule-associated protein 2 (MAP2) (**Fig. 1C**). Maximal neurite-like outgrowth was reached after a treatment period of ~24 hours (**Fig. S1**), and Y-27632-treated cells consistently exhibited TuJ1- and MAP2-positive cell projections in three different (transcriptionally-heterogeneous) GBM cell models (**Fig 1D**).

**Figure 1.**
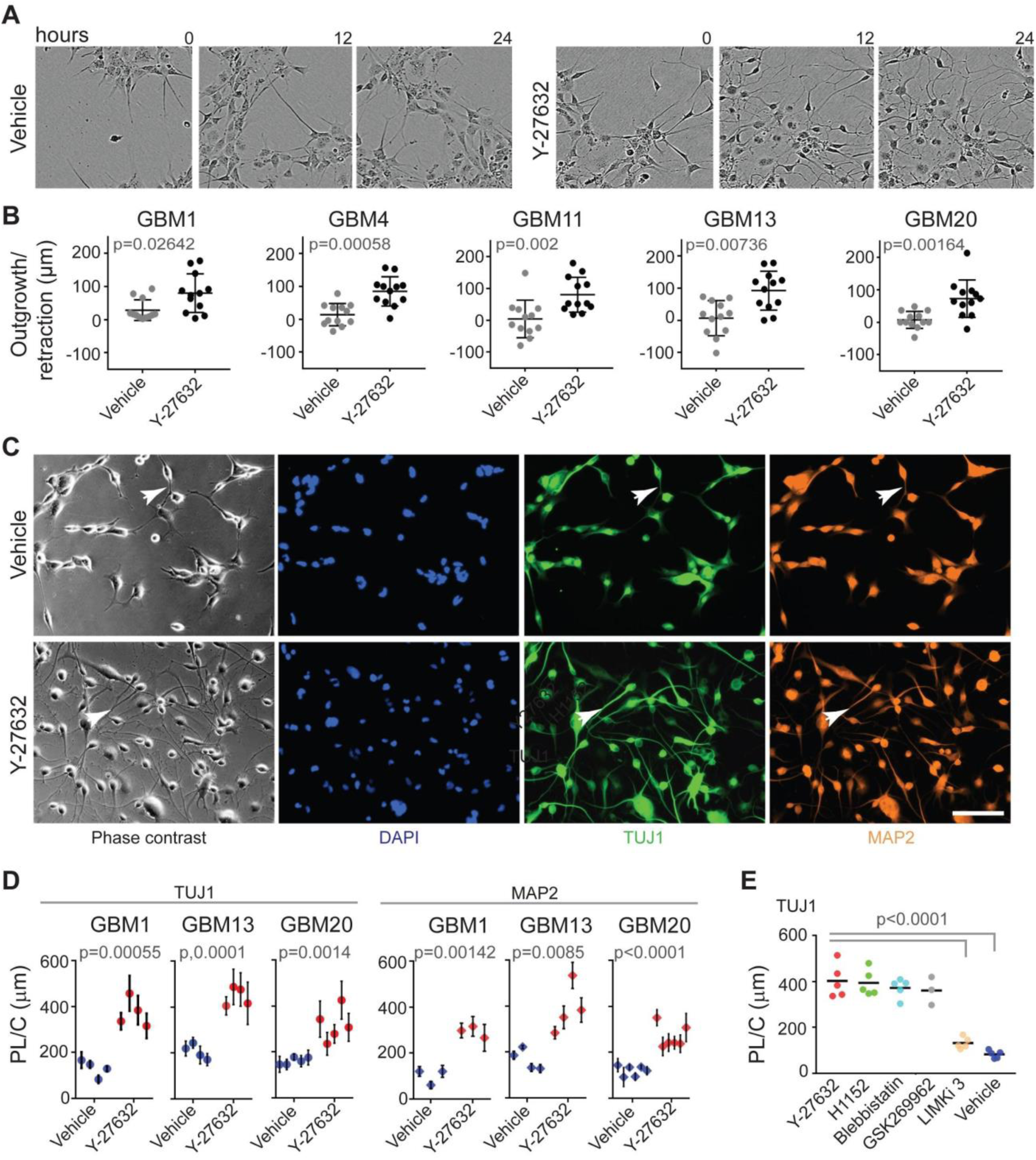
Chemical ROCK inhibition induces neurite-like projection outgrowth in patient-derived GBM cell models. (A) Time-lapse microscopy still-frame images of GBM1 cells treated with vehicle (H_2_O) or Y-27632 (20 μM) at the indicated time points. (B) Projection outgrowth and retraction (μm) per individual cell (represented by a dot) determined by t_o_ versus 24-hour time point comparisons using time lapse microscopy for the indicated GBM lines treated with vehicle (H_2_O) or Y-27632 (20 μM). Plots indicate the median with 95% confidence intervals. (C) Immunocytochemistry images of projections in GBM1 cells (white arrowheads) stained for the neuron-specific microtubule-associated protein 2 (MAP2) and class III β-tubulin (TuJ1). Scale bar, 50 μm. (D) Y-27632-induced TUJ1- and MAP2-positive neurite-like projection length per cell (PL/C, 24 hours) in the indicated cell models compared to the vehicle control. Biological replicate (dots) are shown as mean ± standard deviation. (E) TuJ1-positive projection length per cell (PL/C, 24 hours) in GBM1 cells treated with vehicle (H_2_O) or the indicated ROCK pathway inhibitors. Biological replicates are shown as dots, black bar depicts the mean. p-values were determined by the Mann-Whitney U test (two-tailed; B), Student’s t-test (2-tailed, equal variance; D), and One-way ANOVA (E).

Next, we tested whether structurally-distinct ROCK inhibitors (H1152 and GSK269962) would recapitulate the neurite-like outgrowth in GBM1 cells. H1152 (10 μM; (Ikenoya et al. 2002)) and GSK269962 (5 μM; (Doe et al. 2007)) resulted in neurite-like projection outgrowth that was comparable to the one induced by Y-27632 (~4-fold, p<0.0001; **Fig 1E**). Moreover, we disrupted two different downstream effectors of ROCK using Blebbistatin, an inhibitor of the nonmuscle myosin II protein (NM II; (Straight et al. 2003)), and LIMK inhibitor 3 (LIMKi3), a suppressor of LIMK-dependent cofilin phosphorylation (Ross-Macdonald et al. 2008). Blebbistatin (2.5 μM) markedly increased neurite-like projection outgrowth in GBM1 cells (p<0.0001), whereas LIMKi3 (10 μM) did not induce this phenotype compared to the control (**Fig 1E**). Taken together, these data indicate a robust chemical induction of neurite-like outgrowth in adherently-cultured GBM cell models that was mediated by the disruption of ROCKi-dependent activation of NM II, rather than LIMK protein.

### A *de novo*-induced network (iNet) topology via ROCKi and GBM cell projection elongation

To determine whether the ROCKi-induced projection elongations provided the basis for an integrated multicellular system, we used mathematical network analysis through mapping GBM1 cell bodies and protrusions under the assumption that they are connected objects, equivalent to nodes and edges in a discrete network (Jackson et al. 2017) (**Fig. 2A**). In contrast to network topologies obtained for control cells (hereinafter referred to as Con), the logical network topology indicators ‘degree and betweenness centrality’ (Newman 2006) were significantly elevated in the Y-27632-treated cultures (**Fig. 2B**), hence indicating a *de novo*-induced cellular network phenotype characterized by an increased range of cell-to-cell connectivity (hereinafter referred to as iNet). Next, we experimentally tested implications of the iNet cell-to-cell connectivity on cellular motility by changing the relative distance between cells using low (<10% confluency) or medium (>50% confluency) cell seeding densities. This assay revealed a marked decrease in iNet cellular motility solely under iNet/cell-to-cell contact-permissive conditions (>50% confluency; p<0.0001; **Fig. 2C**). Moreover, the iNet cells retracted their projections within eight hours of ROCK inhibitor withdrawal, demonstrating reversion of the cell projection-based ‘web’ phenotype (hereinafter referred to as Rev) (**Fig. 2D, Figure S2A and B**). Time lapse microscopy, Ki67-staining, and clonal cell growth analysis indicated comparable proliferation capacities of GBM1 cells under Con, iNet, and Rev phenotypic conditions (**Fig. 2E, Figure S2C**). In addition, RNA-seq transcriptional comparison of iNet and Rev GBM1 cellular profiles revealed Adhesion Molecule With Ig Like Domain 3 (AMIGO3), an inhibitor of axonal growth (Ahmed et al. 2013), as the most downregulated iNet-associated transcript (**Fig. 2F**). Further analysis using The Database for Annotation, Visualization and Integrated Discovery (DAVID; (Huang et al. 2009b; Huang et al. 2009a) and Gene Set Enrichment Analysis (GSEA; (Subramanian et al. 2005)) suggested iNet-specific enrichment of axonal guidance-regulating genes (**Fig. S3A and B**), and positive iNet signature correlations with cell projection growth, cytoskeletal organization, cell cycle progression, and anti-apoptotic pathways (**Fig. 2G**).

**Figure 2.**
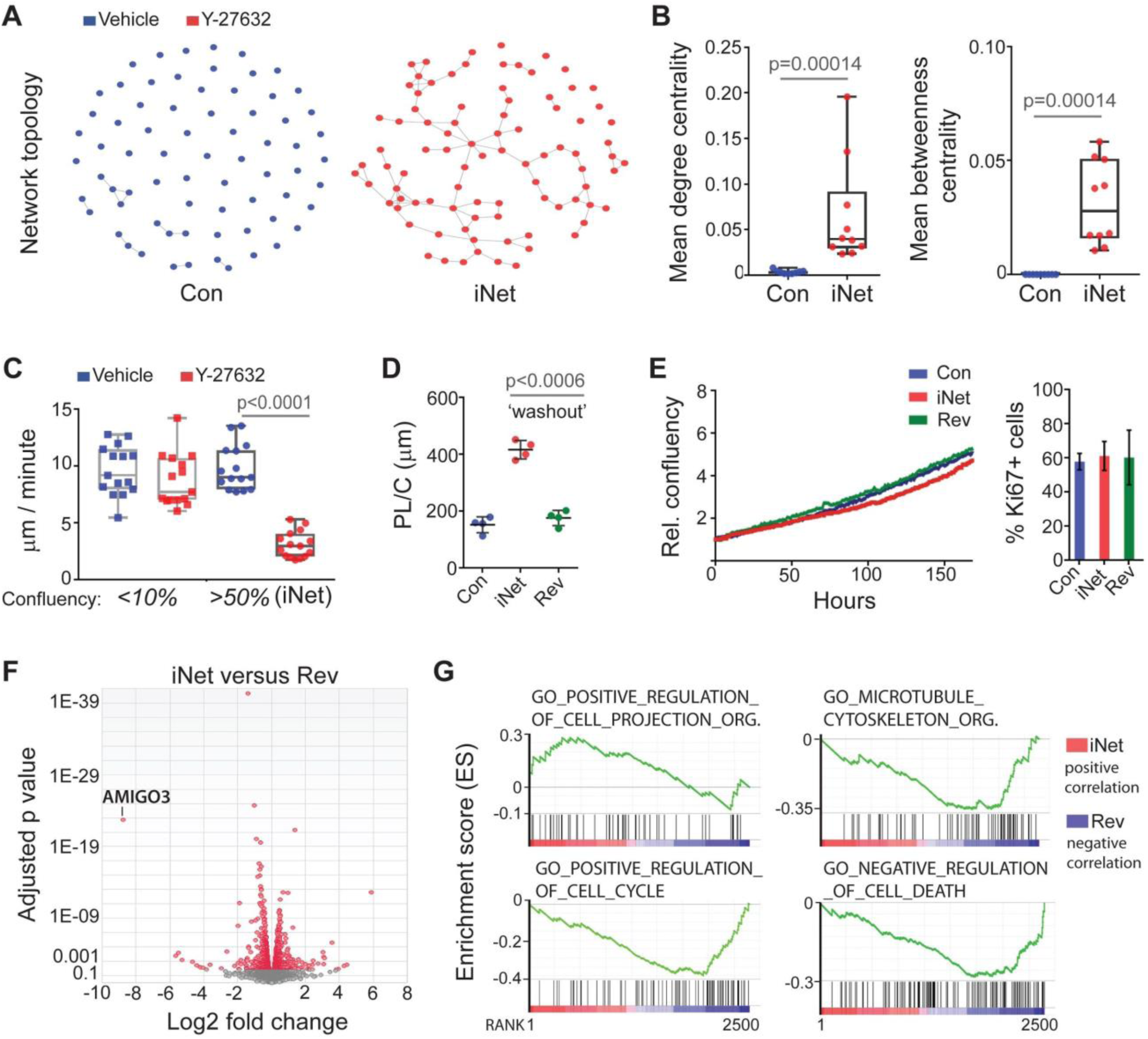
Chemically-induced cell projection outgrowth enables de novo network formation in GBM cells. (A) Representative topologies of control (Con; left) or *de novo*-induced GBM1 cell networks (iNet, right) in presence of vehicle (H_2_O) or Y-27632 (20 μM), respectively. (B) Con versus iNet network modal degree (left) and betweenness (right) values (biological replicates). (C) Individual GBM1 cell movement distances (dots) under low confluency versus iNet-permissive (>50% confluency) culture conditions in the presence of vehicle (H_2_O) or Y-27632 (20 μM). (D) Neurite-like projection length per GBM1 cell (PL/C) in Con, iNet, and Rev (Y-27632 ‘washout’) conditions. (E) Left, real-time assessment of GBM1 cell growth (confluency shown as fold change normalized to t0 values) in Con, iNet, or Rev (Y-27632 ‘washout’) conditions. One out of three biological replicates (see Supplementary File 1) is shown. Right, percentage of cells expressing Ki67 protein in Con, iNet, or Rev conditions. (F) Volcano plot comparing transcriptional RNA-seq profiles of iNet and Rev phenotypes in GBM1 cells. (G) GSEA plots comparing transcriptional iNet and Rev profile correlations with the indicated GO pathways. The p-values were determined by the Mann-Whitney U test (one-tailed; B), and One-way ANOVA (C, D).

### The iNet phenotype increases GBM multicellular co-operation

To test whether the growth of GBM1 cell projections enabled multicellular co-operation, we assessed calcium and fluorescent dye transmission after establishing the ROCKi-induced iNet phenotype (**Fig. 3A and B**). Upon focused laser stimulation, the Fluo-3 (Minta et al. 1989) signal intensity increased in the neighboring GBM1 cells, and maximum intensities were markedly elevated under iNet compared to Con phenotypic conditions (p=0.012; **Fig. 3C**). Less than 40% of Con cells participated in the Fluo-3 transmission waves with Fluo-3 signal angles ranging between 60 and 320 degrees (**Fig. 3D**). In contrast, ~80% of iNet cells participated in the calcium signal extensions exhibiting angles of >320 degrees (**Fig. 3D**). This long range cell-to-cell communication was corroborated by scrape-loading dye transfer experiments (el-Fouly et al. 1987) showing a significant increase in the cell-to-cell transfer of Lucifer yellow in iNet compared to Con cells (>2-fold, p=0.005; **Fig. 3E**), thus indicating gap junction-mediated cell-to-cell communication. Furthermore, confocal time lapse imaging revealed significant trafficking of lysosomes and mitochondria between the iNet cells, which was limited by the lack of cell projection elongations in Con cells (**Fig. 3F and G, Videos S11-14**).

**Figure 3.**
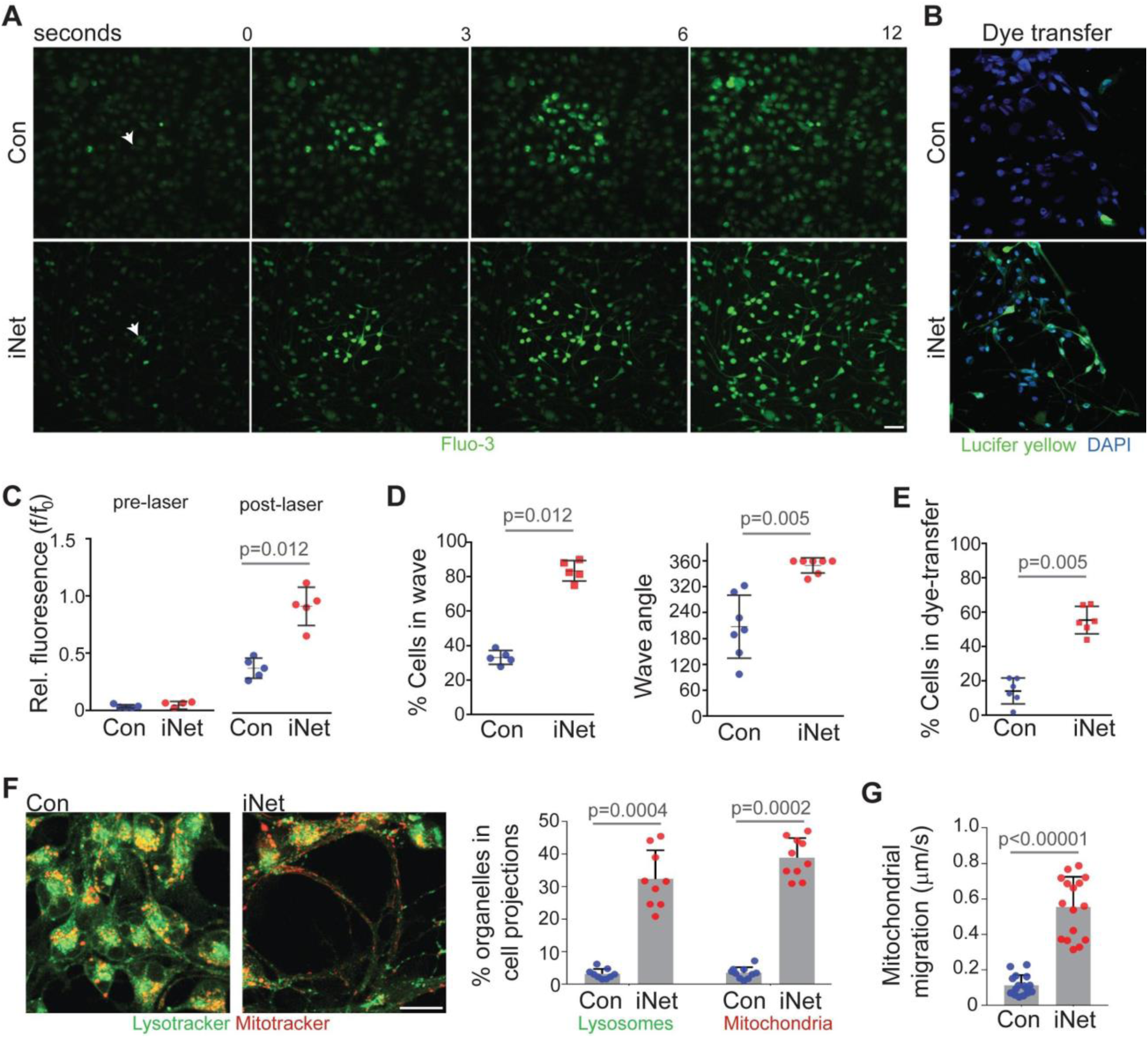
The iNet phenotype is characterized by GBM multicellular co-operation. (A) Still frame time lapse microscopy images of intracellular calcium wave transmission upon laser irradiation of a single GBM1 cell (arrowhead) in Con or iNet conditions. Scale bar, 30 μm. (B) Microscopic false color image of lucifer yellow dye transfer using scrape loading technique comparing Con with iNet conditions. (C) Quantification of calcium signal wave intensity (relative fluorescence) pre- and post-laser irradiation in Con or iNet phenotypic conditions. (D) Percentage of total Con or iNet cells transmitting the calcium wave (left), and wave transmission angle (right). Values (dots) depict biological replicates. (E) Quantification of Con or iNet Lucifer yellow-transferring cells per biological replicate (dots). (F) Left, confocal microscopy imaging of indicated organelles. Scale bar, 20 μm. Right, quantification of lysosomal and mitochondrial localization in Con versus iNet cell projections. (G) Migration distances of individually-traced mitochondria in Con or iNet phenotypic conditions. p-values were determined by the Mann-Whitney U Test for all comparisons.

### The iNet phenotype promotes proliferation and survival of GBM cells post radiation

As multicellular brain tumor networking has been linked to pro-tumorigenic functions (Osswald et al. 2015), we examined potential ionizing radiation resistance-promoting effects of the iNET phenotype in GBM1 cells. Two hours after 0, 2, 8, or 20 Gy radiation, a dose-dependent increase of DNA double-strand breaks was detected by □H2AX staining (Kuo and Yang 2008), and similar DNA damage levels were observed in Con and iNet GBM1 cells (**Figure S4A**). However, only the iNet phenotype was associated with a marked increase in cellular projection outgrowth (**Figure S4B**) as well as significantly higher proliferation rates 5 days after 8 and 20 Gy radiation when compared to Con and Rev (p<0.0272; **Figure 4A**). These results were corroborated by cell fate tracing of 15 different Con, Rev, and iNet cells after 0, 2, 8, and 20 Gy radiation. Most iNet cells were connected via cell projections and displayed a low motility phenotype (**Figure S5**). Taking the total of 45 analyzed cells in all radiation treatments into account, both cell cycle arrest and cell death were reduced by ~90%, and multinucleation by ~80% in iNet cells compared to their Con and Rev counterparts. Thirty-five irradiated iNet cells underwent mitosis, whereas only 7 and 8 Con and Rev cells divided, respectively (**Figure S5**).

**Figure 4.**
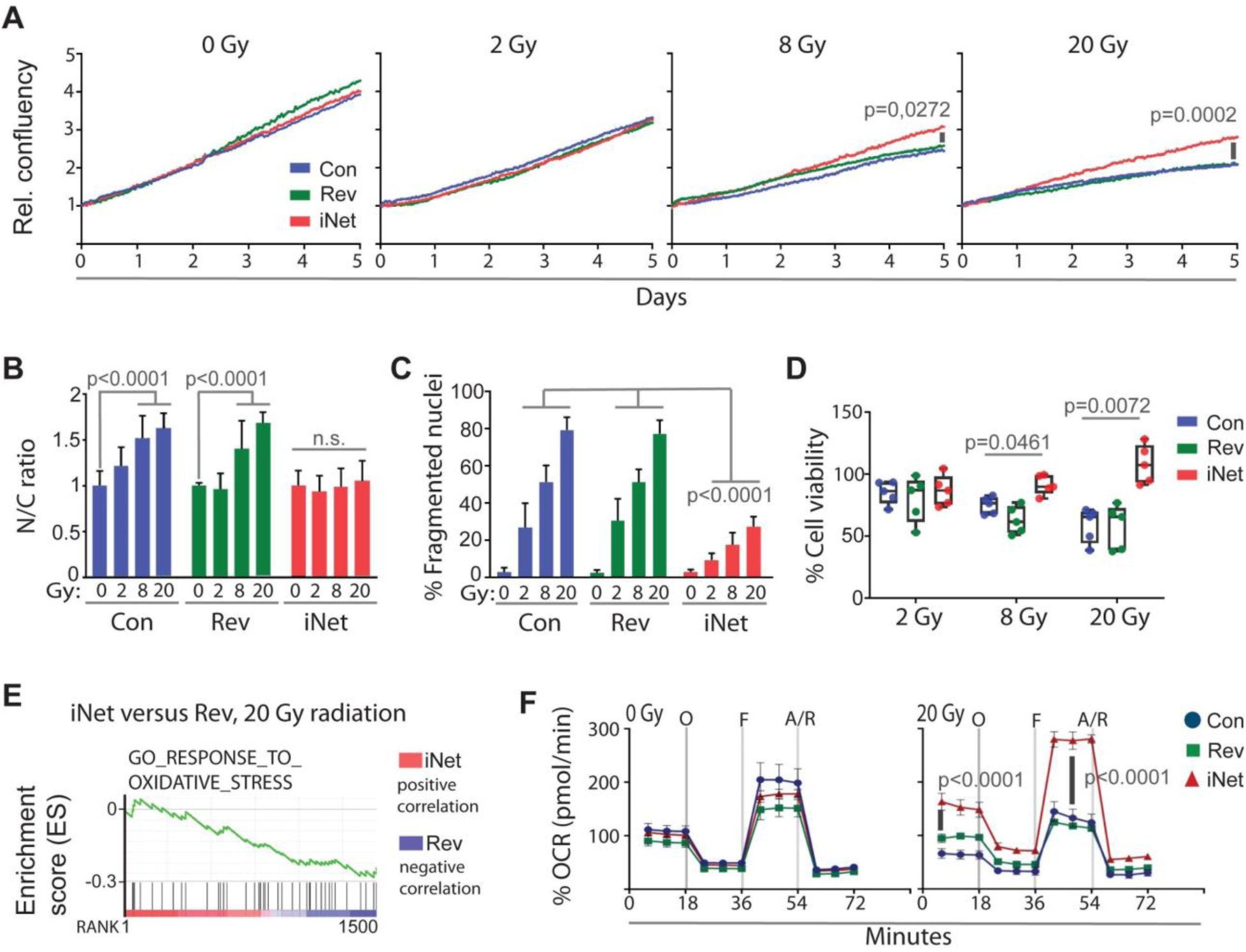
GBM cells connected by iNet phenotype sustain their growth upon radiation treatment. (A) Real-time assessment of GBM1 cell growth (confluency shown as fold change normalized to t_0_ values) in Con, iNet, or Rev phenotypic conditions upon radiation treatment with the indicated doses. One out of three biological replicates (see Supplementary File 1) is shown and p-values were determined by 2-way Anova. Chemically-induced multicellular networks promote GBM cell stress resistance. (B) GBM1 nuclear (N) to cytoplasmic (C) ratio in Con, Rev, and iNet phenotypic conditions 5 days after irradiation with the indicated doses (Gy). (C) Quantification of fragmented nuclei (%) in Con, Rev, or iNet GBM1 cells 5 days after irradiation with the indicated doses (Gy). (D) Biological replicates (dots) of cell viability measurements in Con, Rev, or iNet GBM1 cells 5 days after irradiation with the indicated doses (Gy), normalized to the non-irradiated control. (E) GSEA plot comparing transcriptional iNet and Con profile correlations with the indicated GO pathway. (F) Mitochondrial bioenergetic (extracellular flux) analysis of GBM1 cells 5 days after 20 Gy irradiation. OCR, oxygen consumption rate; O, Oligomycin; F, FCCP; A/R, Antimycin/Rotenone. Data are three biological replicates and error bars depict the standard deviation of the mean. The p-values were determined by One-way ANOVA (A, C, E), and Two-way ANOVA (b) tests.

To further compare radiation-induced damage in GBM1 cells under Con, Rev, and iNet conditions, we quantified cytoplasmic swelling and nuclear fragmentation. The nuclear-to-cytoplasmic ratio remained unchanged in the iNet cells, whereas a dose-dependent increase in cytoplasmic area expansion was observed in their Con and Rev counterparts (**Fig. 4B**). Consistently, the number of GBM1 cells exhibiting nuclear fragmentation 5 days after radiation treatments (2, 8, and 20 Gy) significantly decreased in iNet compared with Con and Rev phenotypic conditions (>2-fold; p<0.001; **Fig. 4C**). Analysis of the iNet cells showed that the vast majority of nuclear fragmentation (≥88%) appeared outside of the multicellular networks and within a cellular subpopulation (<5% of total cells) that remained unconnected (**Fig. S6**). Concomitant luminescent cell viability assays indicated a marked sustainability of iNet cell survival/proliferation after 8 and 20 Gy radiation doses (p<0.0461). In agreement with glioma stem cell-associated radioresistance (Bao et al. 2006), only high-dose radiation (20 Gy) reduced GBM1 cell survival in Con and Rev conditions by approximately 50%, whereas the iNET cells remained viable (p=0.0072; **Fig. 4D**). Accordingly, we carried out RNA-seq transcriptional profiling and GSEA upon 20 Gy radiation comparing Rev and iNet mRNA expression signatures. The iNet profile was positively-correlated with ‘mitotic cell cycle’, ‘negative regulation of cell death’, ‘P53’, ‘hypoxia’, and ‘oxidative stress’ pathways (**Fig. 4E, Fig. S7**). Consistent with the latter being an expected consequence of ionizing radiation (Azzam et al. 2012), extracellular flux analysis 5 days after high-dose radiation (20 Gy) showed that basal oxygen consumption rate (OCR), and OCR following mitochondrial stress test, were significantly elevated in iNet, compared to Con and Rev conditions. This indicates an iNet phenotype-mediated ability of GBM cells to maintain their oxidative metabolism under radiation-induced stress, which has been shown to be required for GBM cellular fitness and viability (Polson et al. 2018).

## DISCUSSION

Our results show that ROCKi significantly promotes the outgrowth of cell projections in a spectrum of GBM subtype stem-like models, which in turn enabled the *de novo* formation of multicellular networks. The ROCKi-induced cell projections exhibited a neurite-like appearance and were immunopositive for the neuronal cell and dendrite marker MAP2, which has been shown to be expressed in both low- and high-grade diffuse brain tumors (Wharton et al. 2002). The observed reversibility of the iNet phenotype even after 5 days of ROCKi treatment, the unaltered proliferation and clonal growth capacity of the ROCK inhibitor-treated GBM cells as well as their RNAseq/GSEA-based profiles were all indicative of dynamic morphological changes rather than cell fate bias/differentiation (Piccirillo et al. 2006; Wurdak et al. 2010; Carén et al. 2015). Consistently, the establishment of the iNet phenotype was characterized by a gene expression profile indicative of cytoskeletal rearrangement, cell cycle progression, and axonal outgrowth pathways, with the latter being previously described as a downstream effect of ROCKi in non-cancerous cells of the central nervous system (Fujita and Yamashita 2014). In agreement with the notion that tumors can function as communicating networks (Winkler and Wick 2018), the induced neurite-like projection ‘web’ provided enhanced GBM cellular connectivity as observed by wide-ranging calcium signal transmission and lysosomal as well as mitochondrial exchange between cells in the network. This iNet activity was associated with a survival and proliferation advantage under radiation treatment that may be explained by the connected cell continuum being able to spread the load of the radiation-induced mitochondrial damage and calcium perturbation, which has been linked to cell death (Orrenius et al. 2003).

Due to their eye pressure-lowering and neuroprotective characteristics, some ROCK inhibitors have been trialed in the clinic as anti-glaucoma (Honjo and Tanihara 2018) and spinal cord injury treatments (NCT00500812). A number of preclinical studies have suggested that ROCK inhibition has therapeutic implications in cancers (Wei et al. 2016; Rath and Olson 2012), hence raising the prospect of clinically-testing ROCK-targeting anti-cancer drugs. However, our findings of ROCKi-induced cellular networks and resultant radiation resistance caution against the use of ROCK-targeted pharmacological inhibitors as anti-brain tumor agents.

The herein described experimental approach to induce a connected cellular network phenotype, that is reversible, may overcome a current limitation in studying network function in poorly-differentiated GBM cells that maintain parental tumor phenotypes during ex vivo expansion (Lee et al. 2006; Pollard et al. 2009; Wurdak et al. 2010). A variety of such patient-derived GBM stem-like models have become available as basic and translational research resources (Xie et al. 2015). Thus, it is plausible that iNet GBM cell phenotypes, either mediated by ROCKi or other cytoskeletal/cell communication-promoting factors, could be widely implemented for the discovery and validation of GBM stem cell radiosensitization targets, for example in the context of high-content chemical and genetic screens (Wurdak 2012). The establishment and reversion of the ROCKi-mediated iNet phenotype may aid in the elucidation of intermittent cytoskeletal dynamics and exchange of signals and organelles in the GBM cellular context. Further in vitro and in vivo investigations are warranted to shed light onto the mechanistic link between ROCKi, GBM cell network morphologies, and tumor-promoting functions.

## MATERIALS AND METHODS

### Cell Culture

The patient-derived GBM cell models were previously established and characterized as described in (Wurdak et al. 2010; Polson et al. 2018). The GBM1 (classical/proneural GBM subtype), GBM4 (mesenchymal GBM subtype), GBM11 (mesenchymal GBM subtype), GBM13 (proneural GBM subtype), and GBM20 (proneural/mesenchymal GBM subtype) cell lines represent a spectrum of transcriptionally-heterogeneous primary and recurrent tumors (Polson et al. 2018). All these cell models maintain stem cell-like characteristics under adherent culture conditions, hence allowing for the rapid quantification of cellular responses to chemical perturbations (Wurdak et al. 2010; Wurdak 2012; King et al. 2017; da Silva et al. 2018; Polson et al. 2018). The cells were adherently propagated on plasticware coated with poly-L-ornithin (5 μg/mL, Sigma, P3655) and laminin (5 μg/mL, Invitrogen, 23017-015). The cells were cultured in Neurobasal medium (Gibco; 21103-049) supplemented with 0.5x B27 (Invitrogen, 17504-044), 0.5x N2 (Invitrogen; 17502-048), recombinant human basic fibroblast growth factor (40 ng/mL, bFGF, Gibco, PHG0024) and epidermal growth factor (40 ng/mL, EGF, R&D systems, 236-EG) Cells were maintained in a humidified incubator with 5% CO2 at 37 °C.

### Small molecule inhibitor treatments

For small molecule inhibitor cell profiling, a seeding density of 20,000 cells per cm^2^ was used and small molecule inhibitor treatments started post cell adhesion (overnight). The ROCK pathway inhibitors Y-27632 (20 μM, Sigma), H1125 (10 μM, Tocris), GSK 269962 (5 μM, Tocris), Blebblestatin (2.5 μM, Tocris) and LIM kinase inhibitor (LIMK3, a kind gift from Michael Olson, Ryerson University; 10 μM) were added for 24 hours. For compound withdrawal (‘washout’) experiments, cells were treated with distinct ROCK pathway inhibitors for the indicated period, followed by the removal of the treatment containing media (washing the cells with PBS), and subsequent addition of media lacking the small molecule inhibitors. Cells were assessed 8 hours post compound ‘washout’.

### Live cell imaging and cell proliferation measurements

Live cell imaging was performed using GBM cells seeded at the density of 20,000 cells per cm^2^ (in a 24-well plate format). Cells were treated with Y-27632 or vehicle (H_2_O) and observed/recorded using the IncuCyte ZOOM® live cell imaging system (Essen Bioscience). Phase contrast images were acquired for all conditions every hour over the indicated periods using a 10X objective. Real time movies and confluency/growth curves data was obtained using the IncuCyte ZOOM® software package.

### Clonal cell growth assay

Cells were seeded at a density of 150 cells/well into 24-well plates and allowed to adhere for 24 hours. The next day, the single cells in each well were identified and counted. Treatments using vehicle (H_2_O), or Y-27632 (20 μM, 24 hours) followed by ‘washout’ (compound withdrawal for 8 hours), or Y-27632 (20 μM, 24 hours) were carried out. On day 3 media was refreshed in the distinct conditions and after 7 days, the number of clonally-derived colonies consisting of >4 cells were determined.

### Immunostaining

Cells were fixed with 4% (w/v) paraformaldehyde (PFA, Sigma-Aldrich-P6148) for 10 minutes and blocking was carried out for 1 hour at room temperature with staining buffer (10% FBS in PBS) supplemented with 0.03% (v/v) Triton X-100 (Sigma-T8787). Next, the cells were incubated at 4°C overnight in staining buffer with the following primary antibodies: anti-neuronal class III β-tubulin (TuJ1, 1:300, Covance, cat# 801202); anti-microtubule-associated protein 2 (MAP2, 1:1000, Abcam, cat# ab5392); anti-Ki67 (1:200; Abcam; ab16667) and anti-phospho-H2AX (γH2AX; 1:800; Merck; JBW130). The cells were then incubated in staining buffer for 1 hour in the dark at room temperature with the following secondary antibodies using a 1:500 dilution: goat anti-mouse IgG with Alexa fluor 488 conjugate (Life technologies, cat# A11029); goat anti-chicken IgG with Alexa fluor 647 conjugate (Life technologies, cat# A21449), and Cy3 (Jackson ImmunoResearch;711-165-152; conjugated). The fluorescence signal was detected using an EVOS digital inverted fluorescence microscope or Nikon A1R confocal microscope.

### Cell projection quantification using NeuriteTracer

Images of neurite-like projection formation were obtained as described above and subject to image analysis using the NeuriteTracer plugin for ImageJ (Fiji) (Pool et al. 2008). Threshold parameters were defined using the vehicle cells. The TUJ1 or MAP2-stained projection outgrowth per cell (PL/C, μm) was calculated as follows: total projection outgrowth length divided by the number of total (DAPI-stained) nuclei.

### Network analysis

Cell connections were manually annotated from immunostained images of control- and Y-27632-treated cells using ImageJ. Two cells were considered connected if their TuJ-positive projections overlapped in the images. This process yielded maps of cell association. These associations were subsequently converted into GraphML format using in-house software (convertGML, available on Github at https://github.com/alastair-droop/associationGML). Per node, the degree and betweenness statistics were generated from the cell association graphML files using the igraph (Csardi and Nepusz 2006) and package in R/numerical analysis (R core team 2018) software. Network measures were calculated using the mean of the individual node statistics for each map. A total of nine Con and ten iNet maps (representing biological replicates) were processed. GraphML files are available online at https://doi.org/10.6084/m9.figshare.7270511.v1.

### Cell movement tracking

Cells were seeded at two distinct confluencies (>10, 7.5 ×104 cells/cm^2^ or <50%, 20×104 cells/cm^2^). Following 24 hour treatment with vehicle (H_2_O) or 20 μM Y-27632, cells were monitored using the IncuCyte ZOOM system. Images were taken every 30 minutes and cellular tracking was performed on 15 distinct cells per condition from the obtained videos tracing the nuclei as the ‘point of movement’ via ImageJ software analysis (http://rsb.info.nih.gov/ii).

### Calcium signaling and Lucifer yellow dye assays

GBM1 cells were treated with vehicle (water) or 20 μM Y-27632 for 24 hours to establish the Con and iNet cellular phenotypes. The following day, Fluo-3, AM (ThermoFisher; F23915) was added to the cells and incubated for 30 minutes in the dark at room temperature. The cells were then washed with PBS and kept in the dark for additional 30 minutes. Live cell imaging was done using the Zeiss LSM780 on a Zeiss Observer Z1 equipped with a Coherent Chameleon laser. Laser injury was performed using 100% of the Coherent Chameleon laser at 800 nm. Images were taken every 1.95 sec. Video analysis was carried out using ImageJ software and calcium peak intensity and the percentage of cells in the wave were determined by the quantification of fluorescence intensity in 10 randomly-chosen cells before and after laser damage. The Fluo-3 transmission wave angle was assessed using the ImageJ ‘angle tool’. The number of cells that transmitted the Fluo-3 signal was determined in 5 independent experimental repeats.

For the dye-loading via scratch assays, the Con and iNet cellular phenotypes were established in GBM1 cells (as described above), cells were rinsed with PBS before the addition of 0.05% lucifer yellow (ThermoFisher Scientific; L1177). A scratch wound was induced on the cells using a pipette tip and the dye was left on the cells for 2 minutes, allowing for its uptake. The cells were then rinsed and with PBS (superfluous dye removal) and incubated with GBM media for 8 minutes. After that period life cell images were taken immediately on the A1R confocal microscope.

### Mitochondria and Lysosome live cell staining and imaging

After establishing Con and iNet phenotypes in GBM1 cells using vehicle (H_2_O) or 20 μM of Y-27632 for 24 hours, the lysosomes of the cells were stained using LysoTracker (50 nM; Thermo Fisher Scientific, L7528), and the mitochondria were stained using MitoTracker (25 nM; Thermo Fisher Scientific, M7512). LysoTracker incubation was for 2 hours, while MitoTracker incubation was for 1 hour. Subsequently, cells were fixed with 4% (w/v) PFA for 10 minutes. Images were obtained using the A1R confocal microscope. For live cell imaging, images were taken every 30 seconds, for a total of 20 minutes, using the Zeiss LSM780 on a Zeiss Observer Z1 microscope.

### Radiation treatment and cellular assays comparing Con, Rev, and iNet phenotypic conditions

For cell irradiation, 20,000 cells per cm^2^ were seeded into 24 well plates and Con, Rev, and iNet phenotypes were established in GBM1 cells via vehicle (H_2_O) exposure, Y-27632 addition (20 μM) for 24 hours followed by ‘washout’ (compound withdrawal for 8 hours), and Y-27632 treatment (20 μM for 24 hours), respectively. Subsequently, cells were irradiated with 0, 2, 8, and 20 Gy using the RadSource RS2000 irradiator

For cell behavior profiling of irradiated Con, Rev, and iNet GBM1 cells, live cell imaging was carried out immediately after radiation treatment using the IncuCyte ZOOM system. Five Con, Rev, and iNet cells were tracked for each experimental condition (0, 2, 8, and 20 Gy) for 3 days in three independent experiments, hence yielding 15 individually-traced cell behaviors per condition. An observation matrix was created by assessing each of the cells against the five following questions: 1) does the monitored cell possess elongated neurite-like projections?, 2) is the monitored cell visually-connected to other cells via neurite-like projections?, 3) is the monitored cell motile (>10 μm/min for Con and Rev cells and >2.5 μm/min for iNet cells)?, 4) is the monitored cell multinucleated?, and 5) what is the observed cell fate for the monitored cell? The first four parameters were created as dichotomous variables (possible outcomes were ‘positive’ or’ negative’). The fifth (cell fate) parameter encompassed the categories: ‘divided’ (cell division occurring during observation period), ‘pre-division’ (no cell division occurring during observation period), ‘cell cycle arrest’ (cells which attempted to undergo mitosis but failed in the attempt), and ‘cell death’ (indicated by cell rounding and detachment). A graphical representation of all observed cell behaviors was generated via heat map.

For the measurement of nuclear/cytoplasmic ratio, regions of interest (ROIs) for the cytoplasmic area were defined by color thresholding using ImageJ (default setting, color space: HSB) and the percentage of area was calculated. Nuclear damage was assessed on cells based on DAPI staining: nuclei presenting with multi- or micro-nucleation segments were considered fragmented.

For the cell viability assay, GBM1 cells were seeded at densities of 10,000 cells per well in white 96-well plates (Greiner bio-one; 655083), and subsequently, Con, Rev, and iNet phenotypes were established (with the aforementioned treatments) and cells were subject to (0, 2, 8, and 20 Gy) radiation treatments. After 5 days, the CellTiter-Glo assay (Promega; G7572) was used, and following equilibration of reagents and cells to room temperature, 100 μL of CellTiter-Glo reagent was added to each and incubation was for 10 minutes in the dark. The luminescence signal was measured using the Mithras LB 940 plate reader.

Extracellular flux-based metabolic assessment of Con, Rev, and iNet GBM1 cells was carried out five days after (0 and 20 Gy) radiation treatments using the Seahorse XF extracellular flux analyzer and protocols described in (Pike Winer and Wu 2014). Cells were trypsinized and seeded into the manufacturer’s microplates at a density of 30,000 cells per well. Before analysis, the medium was replaced with XF base media (Seahorse Bioscience; 102353-100) and transferred to a 37°C non-CO_2_ humidified incubator. For the “Mito stress test”, XF base medium was supplemented with glucose (25 mM; Sigma; G8769), sodium pyruvate (0.5 mM; Sigma; S8636) and L-glutamine (2 mM; Gibco; 35050-061). The final medium was adjusted to pH 7.4 and filtered using a 0.2 μM filter. Oligomycin (1 μM), FCCP (0.5 μM), antimycin and rotenone (0.5 μM) were injected according to the “Mito Stress Test” (Seahorse Bioscience; 101848-400) protocol and results were analyzed using ‘Mito Stress Test’ kit protocols.

### RNA-seq transcriptional profiling

RNA sequencing of GBM1 cells was performed under Rev and iNet conditions. Following a 24-hour treatment with Y-27632 and 8 hour washout, cells were subjected to 0 and 20 Gy. Five days following irradiation, total RNA was isolated from three biological replicates of 0 and 20 Gy/Rev and iNet conditions using RNeasy Mini kits following the manufacturer’s instructions (Qiagen; 74106). The integrity and concentration of RNA were determined using Qubit® RNA Assay Kit in Qubit® 2.0 Fluorometer (Life Technologies, CA, USA). A total amount of 3 μg RNA was used per condition for RNA sample preparation. Sequencing libraries were generated using NEBNext® Ultra^™^ RNA Library Prep Kit for Illumina® (NEB, USA) following manufacturer’s recommendations and index codes were added to attribute sequences to each sample. The clustering of the index-coded samples was performed on a cBot Cluster Generation System using HiSeq PE Cluster Kit cBot-HS (Illumina) according to the manufacturer’s instructions. After cluster generation, the library preparations were sequenced on an Illumina Hiseq platform and 125 bp/150 bp paired-end reads were generated. Sequenced reads were quality-processed, aligned and differential expression analysis performed as described in (Conway et al. 2015), with the exception that the alignment was to the hg38 human reference genome using STAR aligner (v2.2.1) (Dobin et al. 2013). Briefly, reads were trimmed to retain quality and adapters removed, aligned to the human genome and quantified prior to paired sample differential gene expression analysis across groups.

## QUANTIFICATION AND STATISTICAL ANALYSIS

Statistical details are provided in the figure legends. Unless otherwise noted, data were analyzed by a two-sided paired Student’s t-test, or 1-way/2-way ANOVA using Excel (Microsoft Office Professional Plus 2013 package) and/or GraphPad Prism software (version 7.05) assuming normal distribution of the data. The Mann-Whitney U test was used to assess data from several dynamic cellular measurements as indicated in the figure legends. Heatmap and volcano plot visualization was done in Excel (Microsoft Office Professional Plus 2013 package). Network degree and betweenness statistics were generated using the igraph (Csardi and Nepusz 2006) and package in R/numerical analysis (R core team 2018) software.

## DATA AND SOFTWARE AVAILABILITY

Network analysis (GraphML) files are available online at https://doi.org/10.6084/m9.figshare.7270511.v1 and the software at https://github.com/alastair-droop/associationGML).

The RNA-seq transcriptional profiling data has been deposited into The European Nucleotide Archive under accession number PRJEB26811 and unique study name ena-STUDY-

## ACKNOWLEDGEMENTS

We thank Mihaela Lorger and Anestis Tsakiridis for critical advice. B. d. S. and H.W. acknowledge support from ‘The Ian Meek PhD studentship’ from the Brain Tumour Research and Support across Yorkshire charity. H.W. acknowledges support from the Medical Research Council (MR/J001171/1), and A.D. acknowledges support from the UKRI Rutherford fellowship (MR/S00386X/1).

## AUTHOR CONTRIBUTIONS

B. d.S., E.S.P., A.D., G.M. and H.W. conceptualized the study. B.d.S., E.S.P., J.W., M.S. performed experiments. B.d.S., E.S.P., A.D., R.K.M., and H.W. analyzed the data. B.d.S. and A.D. performed network analysis. A.D. and L.F.S. performed bioinformatic analysis and data curation. R. K.M. and S.C.S. provided technical advice. B.d.S. and H.W. wrote the paper. B.d.S., R.K.M., S.C.S., and G.M. edited, and B.d.S., R.K.M., and H.W. revised the paper. H.W. supervised the study.

## DECLARATION OF INTEREST

The authors declare no conflict of interest.

